# Diet-responsive Transcriptional Regulation of Insulin in a Single Neuron Controls Systemic Metabolism

**DOI:** 10.1101/751339

**Authors:** Ava Handley, Qiuli Wu, Tessa Sherry, Roger Pocock

## Abstract

To maintain metabolic homeostasis, the nervous system must adapt and respond to an ever-changing environment. Transcription factors are key drivers of this adaptation, eliciting gene expression changes that can alter neuronal activity. Here we show in *Caenorhabditis elegans* that the terminal selector transcription factor ETS-5 not only establishes the identity of the BAG sensory neurons, but is re-purposed to shape the functional output of the BAG neurons post-mitotically. We find that ETS-5 directly regulates the expression of INS-1, an insulin-like peptide, in the BAG sensory neurons. INS-1 expression in the BAG neurons, and not in other INS-1-expressing neurons, decreases intestinal lipid levels and promotes foraging behaviour. Using *in vivo* analysis, we show that elevated intestinal lipid stores, driven by a high glucose diet, downregulates ETS-5-driven expression of INS-1. Together, our data reveal an inter-tissue regulatory loop by which a single neuron can control systemic metabolism, and that the activity of this neuron is modulated by the metabolic state of the organism.

## INTRODUCTION

Neurons are terminally differentiated, yet retain a degree of plasticity to facilitate responses to ephemeral environments and internal metabolic states (Bhattacharya et al., 2019; Cho et al., 2016; Delaney et al., 2017; Gruner et al., 2014; Pocock and Hobert, 2010; Witham et al., 2016). Neuronal plasticity can be mediated through transcriptional changes that alter synapse formation, receptor composition, and signalling molecules (Bayer and Hobert, 2018; Gruner et al., 2016). In this way, neuronal activity is not necessarily a binary ON/OFF output – but can be modulated depending on the physiological context. Elucidating the mechanisms driving plasticity in neurons that systemically control metabolism will enable us to better understand how organisms form nuanced responses to metabolic needs.

As in other organisms, *Caenorhabditis elegans* integrates diverse sensory inputs to formulate complex behavioural and metabolic decisions that best ensure survival and reproduction (reviewed in (Allen et al., 2015)). A leading example of this integrative process is how *C. elegans* spontaneously switch between three broad activity states: roaming, dwelling and quiescence (Ben Arous et al., 2009; Fujiwara et al., 2002; Hills et al., 2004; Shtonda and Avery, 2006). Roaming is defined by expansive forward movement, whereas dwelling is characterised by multiple pauses and reversals within a restricted area. Despite these differences, both roaming and dwelling are active feeding states. In contrast, quiescent animals are in a non-feeding, sleep-like state that indicates satiety – comparable to mammalian post-prandial somnolence (Fujiwara et al., 2002; Raizen et al., 2008; Shtonda and Avery, 2006; You et al., 2008). These behaviours are ideal readouts of the animal’s internal nutritional state, as they are modulated by intestinal fat storage and food availability (Ben Arous et al., 2009; Hyun et al., 2016; Shtonda and Avery, 2006). Neuronal signalling pathways, such as serotonin, TGF-β, pigment-dispersing factor and insulin-like signalling, act in concert to simultaneously regulate metabolism, foraging behaviour, reproduction and developmental decisions (Ben Arous et al., 2009; Flavell et al., 2013; Palamiuc et al., 2017; Song and Avery, 2012; You et al., 2008) This integrative neuronal control enables the animal to appropriately modulate behaviour and metabolism in response to diverse and dynamic inputs.

While we know that the nervous system can directly influence intestinal fat metabolism, which in turn modifies neuronal functions, our understanding of how the nervous system adapts to changes in nutrient availability, and the molecular mechanisms driving those adaptations is far from complete. One means by which neurons can modify their activity is by altering the types and amounts of signalling molecules they produce, through changes in gene expression. For example, starvation induces transcriptional changes to chemoreceptor and insulin/IGF-receptor expression in the nervous system (Gruner et al., 2016; Gruner et al., 2014; Kimura et al., 2011), and sensory inputs can modulate neuropeptide expression (Rojo Romanos et al., 2017). Currently, however, there is a dearth of knowledge about the transcription factors that define and drive neuronal adaption to nutrient availability. And we are only just beginning to appreciate the extent to which the transcription factors that specify neuronal fates are re-purposed, post-mitotically, to drive adaptive changes in neuronal gene expression (Bhattacharya et al., 2019).

Here, we show that a conserved transcription factor, ETS-5, adaptively regulates an insulin-like peptide in response to a high-glucose diet. ETS-5 is an E-TwentySix-domain transcription factor, homologous to Pet1 in mammals, that is expressed in a specific set of glutamatergic neurons, including BAG and ASG (Brandt et al., 2019; Guillermin et al., 2011). The BAG neurons are well known for their gas-sensing functions – sensing down-steps in O_2_ and acute CO_2_ responses (Bretscher et al., 2008; Hallem and Sternberg, 2008; Zimmer et al., 2009). In parallel to these gas-sensing roles, signals from the BAG neurons control egg-laying, and can modulate the activity of URX neurons to control lipid metabolism (Hussey et al., 2018; Ringstad and Horvitz, 2008). ETS-5 plays a fundamental role in transcriptionally regulating the receptors and neuropeptides involved in these BAG-specific functions (Brandt et al., 2012; Guillermin et al., 2011).

We previously found that ETS-5 regulates intestinal lipid stores and foraging/exploration behaviour through neuropeptide signalling from the BAG and ASG neurons (Juozaityte et al., 2017). Here, we aimed to identify the specific neuropeptide(s) that are controlled by ETS-5 to direct neuron-intestinal communication. In addition, as the BAG neurons are sensory neurons, we examined whether ETS-5 within the BAG neurons are responsive to dietary changes, enabling systemic adaptation to metabolic control. We find that the insulin orthologue INS-1 is directly regulated by ETS-5, and controls intestinal fat levels specifically from the BAG neurons. Remarkably, we find that *ins-1* expression within the BAG neurons is dynamically regulated by ETS-5 in response to a high-glucose diet. Together, our study reveals a mechanism whereby a terminal selector transcription factor modulates insulin expression in the post-mitotic BAG neurons in response to changes to nutritional status. This mechanism enables metabolic and behavioural responses that are flexible to changes in the environment.

## RESULTS

### INS-1 Regulates Intestinal Fat Levels and Foraging Behaviour

Neuropeptide release from the BAG neurons drives foraging behaviour (Figure 1A, (Juozaityte et al., 2017)). However, the identity of these neuropeptide(s) and how they are regulated is unclear. Animals defective for BAG neuropeptide release phenocopy those lacking the BAG terminal selector transcription factor, ETS-5. Therefore, we hypothesized that ETS-5 controls foraging through neuropeptide regulation. To identify the neuropeptide signals that could be regulated by ETS-5, we screened known BAG-expressed neuropeptide mutants for defective exploration behaviour (Figure 1B). Of the neuropeptide mutants tested, we found that *ins-1(nj32*) mutant animals exhibit decreased exploration (Figure 1B). An independently-derived *ins-1(tm1888)* deletion allele also presented defective exploration behaviour (Figure 1C). To verify the exploration defect was specifically due to loss of *ins-1*, we resupplied the *ins-1* genomic region using a fosmid in the *ins-1(nj32)* mutant and found that the exploration defect was significantly rescued (Figure 1C). Therefore, *ins-1* activity is necessary for correct exploration behaviour.

**Figure 1.**
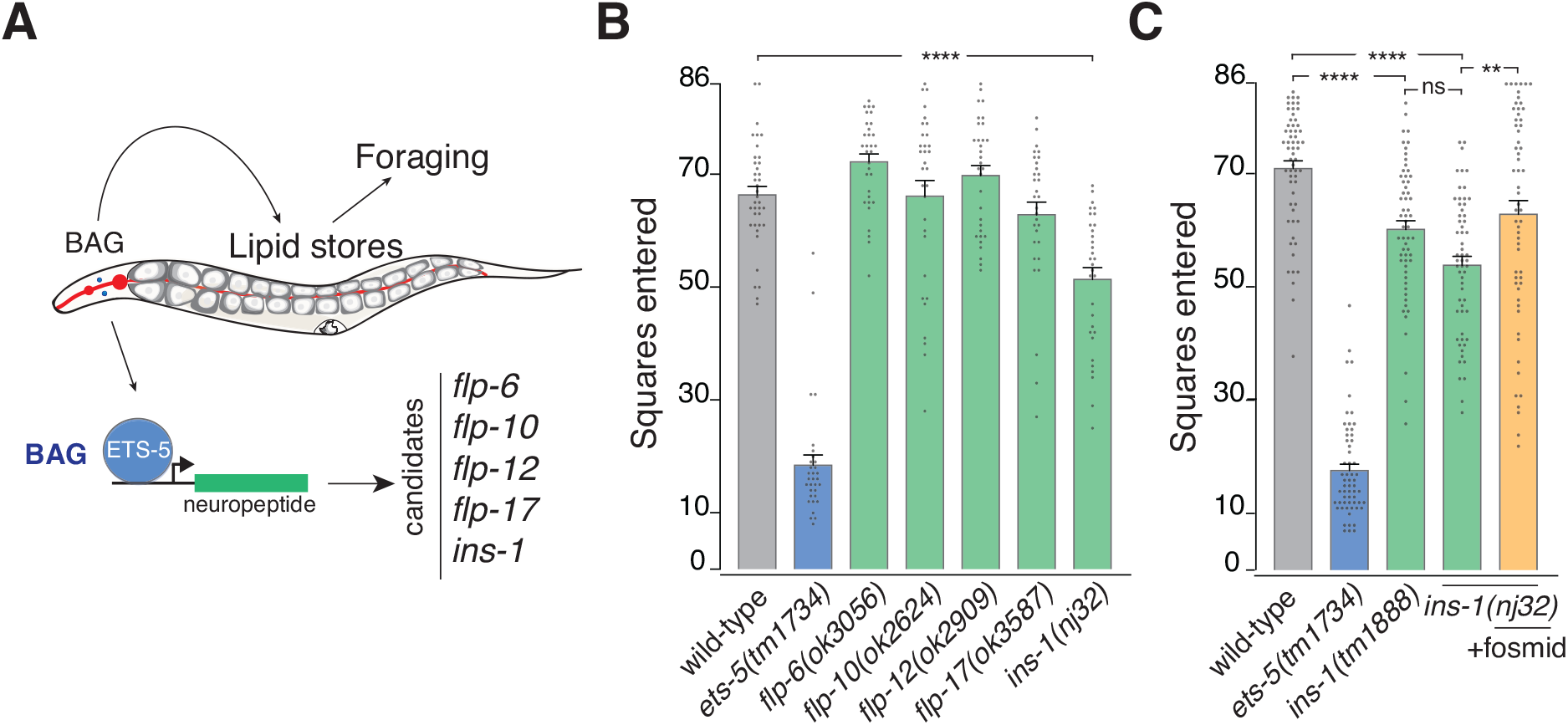
INS-1 controls exploration behaviour in *C. elegans*. (A) ETS-5 acts within the BAG neurons to control intestinal lipid stores via neuropeptide signalling. ETS-5 may regulate any of several neuropeptide genes, listed on the right, known to be expressed within these neurons. (B) Exploration assay data of neuropeptide mutant screen. Data presented as individual worm (points) with 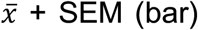, n>30. ****p<0.0001 (one-way ANOVA with Tukey’s correction). (C) Exploration assay in two *ins-1* alleles *tm1888* and *nj32*, with *ins-1(nj32)* + fosmid rescue. Data presented as individual worm (points) with 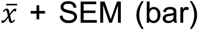, n=59. ****p<0.0001, **p<0.01, ns – not significant (one-way ANOVA with Tukey’s correction).

Our previous work showed that the exploration defect observed in *ets-5* mutant animals was due to increased fat storage (Juozaityte et al., 2017). Therefore, we measured the fat content of each *ins-1* mutant strain by Oil Red O staining and found they store over 30% more fat than wild-type worms (Figure 2A-B). Resupplying the *ins-1* genomic locus to *ins-1(nj32)* animals reduced the fat storage to wild-type levels (Figure 2A-B). To determine whether the increased fat levels observed in the *ins-1* mutant contribute to the exploration defect we used RNA-mediated interference (RNAi) to reduce the levels of the acetyl-coA carboxylase POD-2 in the *ins-1(nj32)* mutant. Reducing POD-2 activity reduces intestinal fat levels by more than 80% (Witham et al., 2016). We found that *pod-2* RNAi rescued the *ins-1(nj32)* exploration defect to wild-type levels (Figure 2C). These data show that, as we observed previously with ETS-5, INS-1 likely modulates exploration behaviour by altering intestinal fat levels. Thus, we hypothesized that INS-1 could be part of the signalling mechanism by which ETS-5 regulates intestinal metabolism. To determine if ETS-5 and INS-1 act in the same genetic pathway to control fat levels we compared Oil Red O staining of the *ets-5; ins-1* double mutant to the respective single mutants (Figure 2D-E). We found that the *ets-5; ins-1* double mutant showed no significant difference in fat content to either single mutant (Figure 2E), suggesting that *ets-5* and *ins-1* act in the same genetic pathway to control intestinal fat levels.

**Figure 2.**
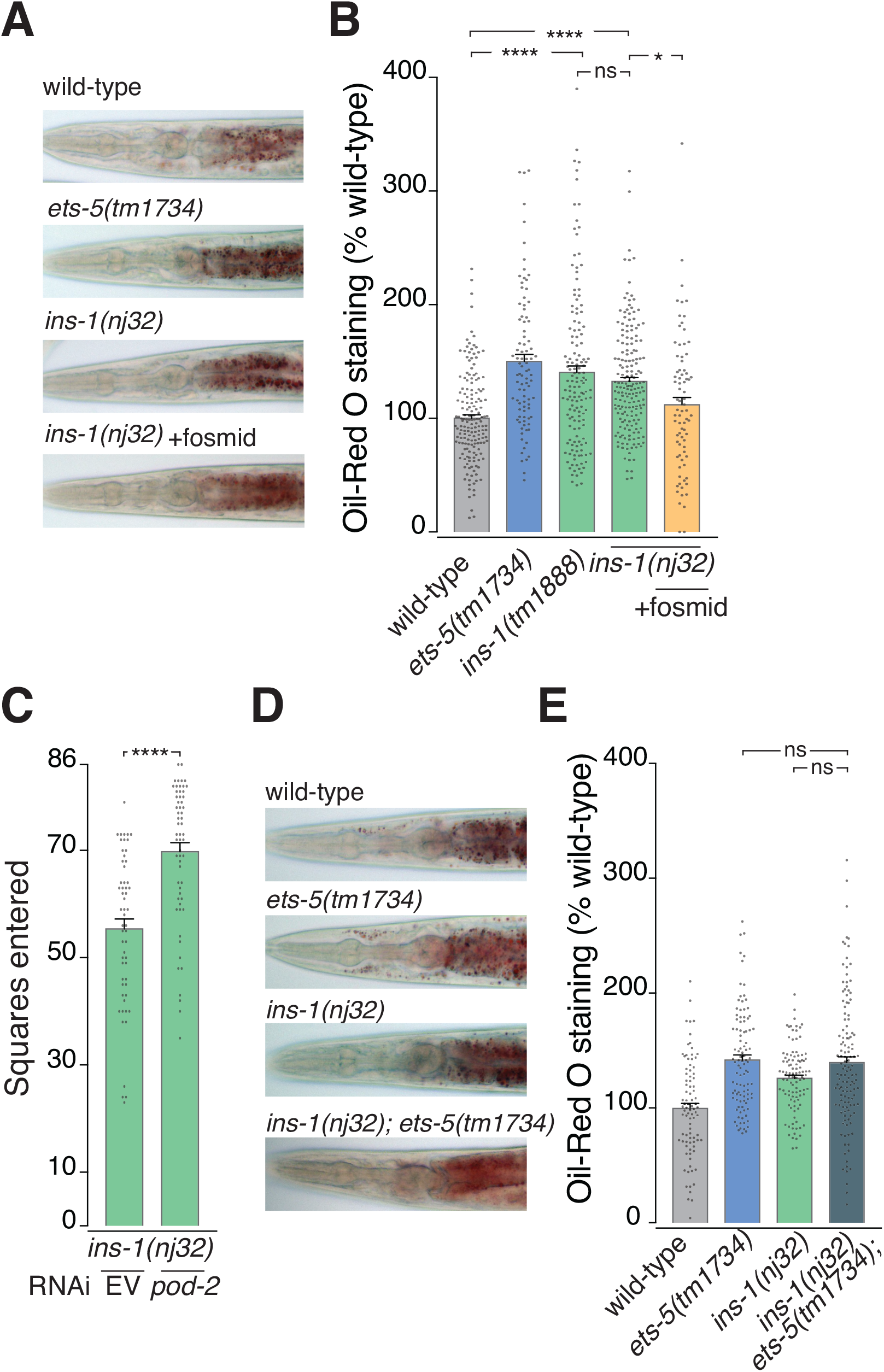
INS-1 regulates intestinal fat levels in the same genetic pathway as ETS-5. (A-B) Oil Red O (ORO) staining image examples (A), and quantification (B) in two *ins-1* alleles *tm1888* and *nj32*, with *ins-1(nj32)* + fosmid rescue. Data presented as individual worm (points) with 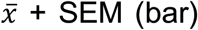, n>80. ****p<0.0001, *p<0.05, ns - not significant (One-way ANOVA with Tukey’s correction). (C) Exploration assay with *pod-2* RNAi, or empty vector (EV) in *ins-1(nj32)*. Data presented as individual worm (points) with 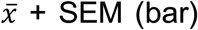, n>50. ****p<0.0001(unpaired t test). (D-E) Double mutant analysis for *ets-5(tm1734)* and *ins-1(nj32)*. Oil Red O staining image examples (D), and quantification of *ins-1(nj32); ets-5(tm1734)* double mutant compared to each single mutant (E). Data presented as individual worm (points) with 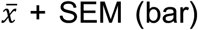, n>80. ns – not significant (One-way ANOVA with Tukey’s correction).

### INS-1 Acts from the BAG Neurons to Control Intestinal Fat Levels

INS-1 was previously shown to be expressed in approximately 10 neurons (Pierce et al., 2001). These include the ASG and BAG neurons - the functionally important sites of ETS-5 for control of fat levels and exploration (Juozaityte et al., 2017). More recently, single cell sequencing analysis reported that *ins-1* is expressed in the BAG neurons and not in the ASG neurons (Cao et al., 2017). To determine if *ins-1* and *ets-5* expression co-localise to the same neurons, we generated transgenic animals carrying a reporter construct that uses a 2.5kb fragment of the *ins-1* promoter to drive expression of nuclear-localised GFP. We then combined this reporter line with a previously generated *ets-5p∷mCherry* reporter strain (Figure 3A) (Juozaityte et al., 2017). The *ins-1p∷NLS-GFP* expression clearly co-localises with *ets-5p∷mCherry* expression in the BAG neurons and not in the ASG neurons (Figure 3A), consistent with recent cell-type specific RNA-seq data (Cao et al., 2017). To demonstrate that the 2.5kb *ins-1* promoter fragment is sufficient to coordinate proper *ins-1* activity, we used this element to drive *ins-1* cDNA expression in *ins-1(nj32)* animals and determined if the exploration defect is rescued. Three independent transgenic lines showed that *ins-1p(2.5kb)∷ins-1* significantly rescues the *ins-1(nj32)* exploration defect (Figure 3B).

**Figure 3.**
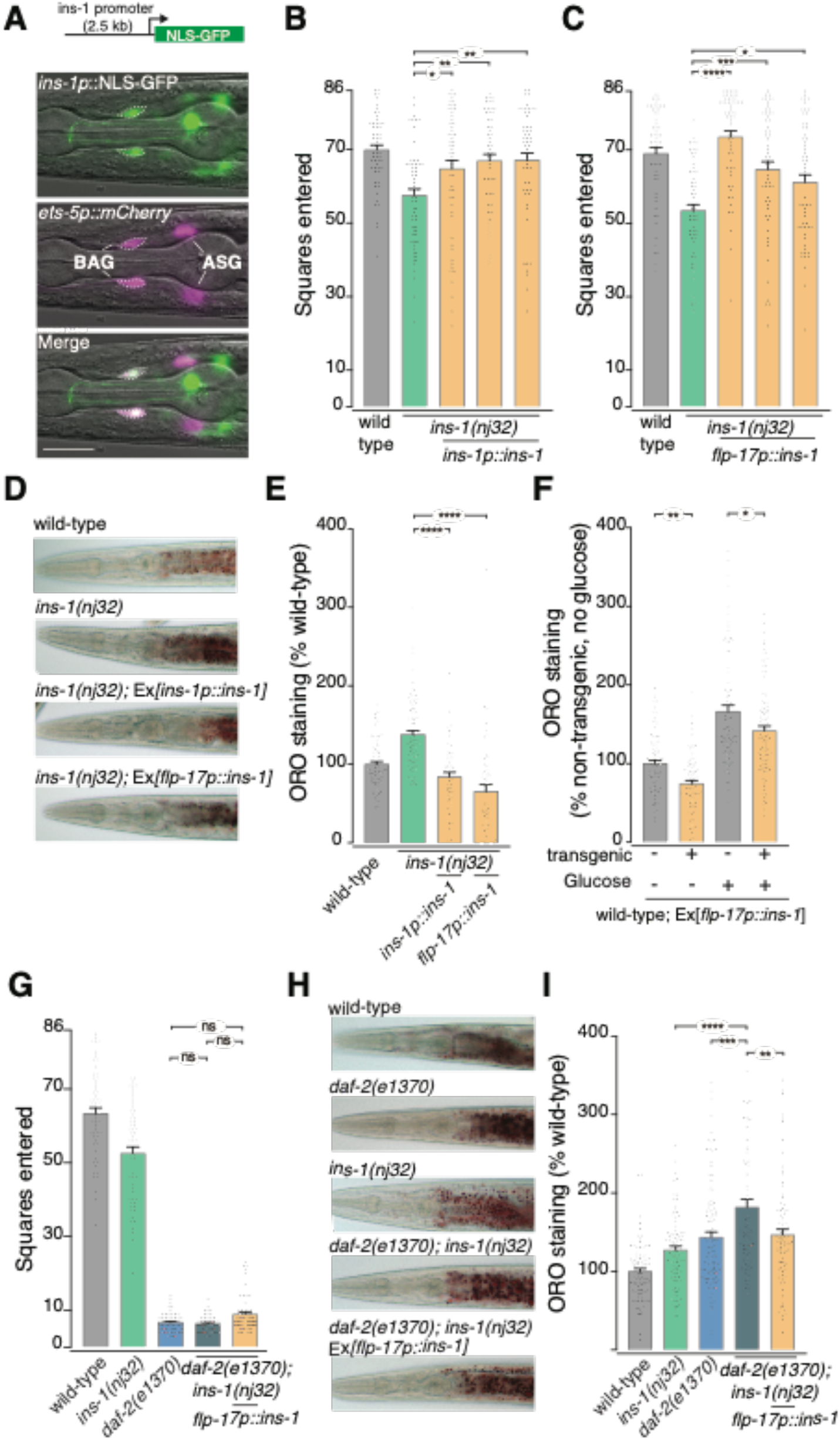
BAG-derived INS-1 regulates intestinal fat levels and exploration behaviour. (A) Co-localisation of *ins-1p∷NLS-GFP* and *ets-5p∷mCherry* reporter gene expression. Scale bar = 50μm. (B-C) Exploration assay data for *ins-1(nj32)* rescue: *ins-1* promoter region driving *ins-1*cDNA (B) and *ins-1*cDNA expression exclusively in BAG neurons using *flp-17* promoter (C). Data presented as individual worm (points) with 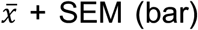, n>50. ****p<0.0001, ***p<0.001, **p<0.01, *p<0.05 (One-way ANOVA with Tukey’s correction). (D-E) Oil Red O images (D) and quantification (E) of *ins-1(nj32)*; *Ex[ins-1p∷ins-1cDNA]* (line 1), *ins-1(nj32); Ex[flp-17p∷ins-1cDNA]* (line 2). Data presented as individual worm (points) with 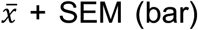, n>50. ****p<0.0001 (One-way ANOVA with Tukey’s correction). (F) Oil Red O quantification of wild-type animals ± *Ex[flp-17p∷ins-1cDNA]* (line1) incubated for 24 hours on OP50 or OP50 + 40mM glucose. Data presented as individual worm (points) with 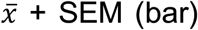, n>70. **p<0.01, *p<0.05 (One-way ANOVA with Tukey’s correction). (G) Exploration assay of *daf-2(e1370); ins-1(nj32)* double mutant in the presence or absence of *Ex[flp-17p∷ins-1cDNA]* (line 2). Data presented as individual worm (points) with 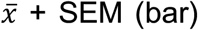, n>50. ns – not significant (One-way ANOVA with Tukey’s correction). (H-I) Oil Red O staining image examples (H) and quantification of *daf-2(e1370); ins-1(nj32)* double mutant in the presence or absence of *Ex[flp-17p∷ins-1cDNA]* (line 2) (I). Data presented as individual worm (points) with 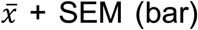, n>70. ****p<0.0001, ***p<0.001, **p<0.01 (One-way ANOVA with Tukey’s correction).

As *ins-1* and *ets-5* expression co-localise only in the BAG neurons, we next asked whether *ins-1* can act specifically from the BAG neurons to alter fat storage and exploration behaviour. To address this question, we used a 200bp fragment of the *flp-17* promoter to drive *ins-1* cDNA expression exclusively in the BAG neurons (Brandt et al., 2012). We found that expressing *ins-1* solely from the BAG neurons rescued the *ins-1(nj32)* exploration defect (Figure 3C). To determine the specificity of the *flp-17p∷ins-1cDNA* action, we mutated the *ins-1* cleavage sites within the *flp-17p∷ins-1* construct that are required for INS-1 to be processed to its active form (Hung et al., 2014). We found that this mutated construct did not rescue the *ins-1(nj32)* exploration defect (Figure S1A-B), showing that proper processing of INS-1 is required to control BAG neuron regulation of exploration, and that no other part of the rescue construct is influencing the *ins-1(nj32)* exploration phenotype. In addition, we replaced *ins-1cDNA* with the coding region of human insulin, and found that human insulin did not rescue the *ins-1(nj32)* exploration defect (Figure S2C).

To determine if the *ins-1* and *flp-17* promoters driving *ins-1* were also sufficient to rescue the increased fat storage in *ins-1(nj32)* animals, we measured Oil Red O staining for one of each rescue line (Figure 3D-E). We found that transgenic *ins-1* expression controlled by the *ins-1* or *flp-17* promoter not only rescued the *ins-1* mutant fat phenotype, but reduced fat levels below that of wild-type animals (~80% of wild-type) (Figure 3D-E). The potency of BAG-derived INS-1 in reducing intestinal fat levels was striking, and prompted us to ask if this decrease in fat levels would persist even if worms were fed a fat-inducing high glucose diet. We crossed a *flp-17p∷ins-1* transgenic line into a wild-type background, then grew animals for 24 hours from the L4 stage on *E. coli* OP50 or OP50 + 40mM glucose. We measured fat levels of these animals, and found that transgenic animals carrying the *flp-17p∷ins-1* array stored significantly less fat than non-transgenic animals on OP50 + 40 mM glucose (Figure 3F). Therefore, BAG-derived INS-1 is capable of potently reducing intestinal fat, even when worms are challenged with a fat-inducing diet.

### INS-1 Regulates Intestinal Fat Levels Independently of the Insulin-like Receptor

We next sought to determine whether BAG-expressed INS-1 acts through the canonical insulin-like signalling pathway. Previous studies have shown that INS-1 can act as an agonist or antagonist of the DAF-2/Insulin-like Growth Factor Receptor depending on the context (Kodama et al., 2006; Pierce et al., 2001; Tomioka et al., 2016). Alternatively, INS-1 may not act through DAF-2 at all, but rather through alternative, unidentified, G-protein coupled receptors (Chalasani et al., 2010). In the context of fat regulation and exploration behaviour, *ins-1* and *daf-2* mutant animals have a similar phenotype - both exhibit increased fat levels and decreased exploration (Ben Arous et al., 2009; Gems et al., 1998; Kimura et al., 1997; McCloskey et al., 2017). Therefore, we hypothesised that BAG-derived INS-1 may act as a DAF-2 agonist to control fat levels and exploration. To determine if INS-1 acts through the DAF-2 receptor, we combined the *ins-1(nj32)* and *daf-2(e1370)* mutants, in the presence or absence of a *flp-17p∷ins-1* rescue line, and measured exploration behaviour and fat levels. In the exploration assay, *daf-2(e1370)* mutant animals explore very little, and the *daf-2(e1370); ins-1(nj32)* double mutant did not show any further reduction in this behaviour (Figure 3G). Further, expressing *ins-1* cDNA in the BAG neurons did not rescue the *daf-2(e1370); ins-1(nj32)* exploration defect (Figure 3G). However, Oil Red O staining showed that the *daf-2(e1370); ins-1(nj32)* double mutant had increased fat storage compared to the *daf-2(e1370)* mutant alone (Figure 3H-I), and this additive effect on fat storage was rescued to *daf-2(e1370)* single mutant levels by resupplying *ins-1* in the BAG neurons (Figure 3H-I). DAF-16, a FOXO transcription factor, is the main downstream target of DAF-2 activity (Lin et al., 1997; Lin et al., 2001; Ogg et al., 1997). We performed exploration assays on *ets-5(tm1734); daf-16(mu86)* and *ins-1(nj32); daf-16(mu86)* double mutants and found that *daf-16(mu86)* explored to a similar extent as wild-type animals, and did not have any effect on either the *ets-5(tm1734)* or *ins-1(nj32)* exploration phenotypes (Figure S2). Taken together, these data suggest that INS-1 does not act through the canonical DAF-2/DAF-16 signalling pathway, and that BAG-derived INS-1 may control intestinal fat levels independently of DAF-2.

### ETS-5 Directly Regulates *ins-1* Expression in the BAG Neurons

The above experiments show that *ins-1* and *ets-5* expression co-localise only in the BAG neurons, that BAG-specific expression of *ins-1* is sufficient for INS-1 function, and that *ins-1* and *ets-5* act in the same genetic pathway to regulate intestinal fat levels. We therefore wanted to determine if ETS-5 regulates *ins-1* expression in the BAG neurons. To address this, we crossed the *ins-1p∷NLS-GFP* reporter strain into the *ets-5(tm1734)* mutant background and measured nuclear GFP expression in the BAG neurons. As the *ins-1* reporter is an extrachromosomal array, we also measured a single ASH neuron in each worm as an internal control for reporter-gene levels (Figure S3). Our data show that *ins-1p∷NLS-GFP* expression in the BAG neurons is significantly decreased in *ets-5(tm1734)* mutant animals (Figure 4A-B). Average GFP expression in the ASH control neuron, where *ets-5* is not expressed, did not significantly change between genotypes (Figure S3B).

**Figure 4.**
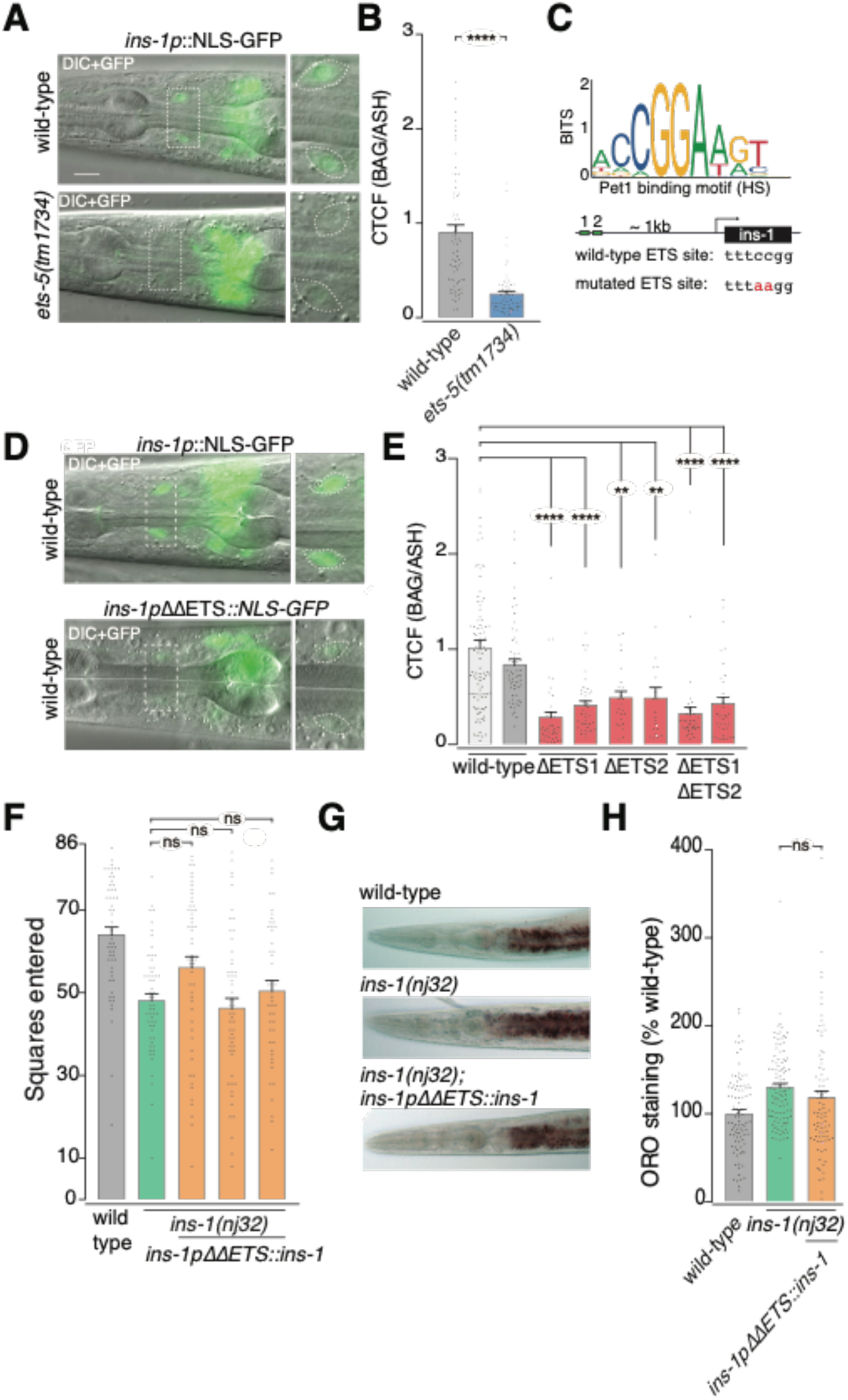
ETS-5 directly regulates *ins-1* expression in the BAG neurons. (A) Representative microscopy images of *ins-1p∷NLS-GFP* expression in *ets-5(tm1734)* compared to wild-type. Scale bar = 10μm. (B) Quantification of GFP intensity (CTCF= calculated total fluorescence) of *ins-1p∷NLS-GFP* in BAG relative to ASH. Data presented as individual nuclei measurements (points) with 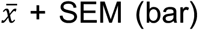, n>70. ****p<0.0001(unpaired t test). (C) Putative ETS/Pet1 binding sites identified, and mutated sequence (red), in the *ins-1* promoter. (D) Representative microscopy images of *ins-1p∷NLS-GFP* with both putative ETS sites mutated (ΔΔETS). (E) Quantification of GFP intensity (CTCF) in BAG relative to ASH, with *ins-1* promoter mutated at ETS1, ETS2 or both, n>20. ****p<0.0001, ***p<0.001, **p<0.01, ns – not significant (one-way ANOVA with Dunnet’s correction). (F) *ins-1(nj32)* exploration rescue using the *ins-1* promoter with both putative ETS sites mutated to drive *ins-1* cDNA. Data presented as individual worm (points) with 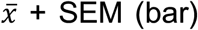, n>45. ns – not significant (One-way ANOVA with Tukey’s correction). (G-H) Oil Red O images (G) and quantification (H) of *ins-1(nj32)*; *Ex[ins-1ΔΔETSp∷ins-1cDNA]* (line 3). Data presented as individual worm (points) with 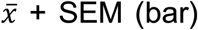, n>50. ns – not significant (One-way ANOVA with Tukey’s correction).

ETS-5 is a key transcriptional regulator of many genes in the BAG neurons, meaning that the downregulation of *ins-1* may be a secondary effect of aberrant ETS-5 activity. To determine if ETS-5 directly regulates the *ins-1* promoter, we searched for the presence of potential ETS binding sites using the Pet1 consensus motif (Khan et al., 2018). We found two such sites in the *ins-1* promoter that are located around −1300 and −1180 from the *ins-1* transcriptional start site (Figure 4C). Mutating the canonical ETS/Pet1 binding motif has been shown to abolish expression of the guanylate cyclase-encoding genes *gcy-31* and *gcy-9* in the BAG neurons (Figure 4C) (Brandt et al., 2012; Guillermin et al., 2011). Two central cytosines form the most highly conserved part of the ETS motif (Wei et al., 2010). We used site-directed mutagenesis to alter these two central cytosine residues to adenine in each ETS motif within the *ins-1p∷NLS-GFP* construct (Figure 4C), which we predict to abolish ETS-5 binding. We generated two independent transgenic lines for each mutagenized construct: ETS site 1 alone, ETS site 2 alone, and both ETS sites mutated. We then measured nuclear-GFP expression in BAG nuclei relative to ASH nuclei. We observed that each ETS site is required for full *ins-1* promoter activity in the BAG neurons (Figure 4D-E). Transgenic lines with both ETS sites mutated (*ins-1pΔΔETS∷NLS∷GFP)* showed no further decrease in promoter activity (Figure 4D-E). These data indicate that both ETS sites within the *ins-1* promoter are required for high expression in the BAG neurons, and that these sites likely act cooperatively, since when one is lost the function of both is abrogated.

Our data show that the ETS binding sites are important for correct expression of the *ins-1* promoter in the BAG neurons. We next determined whether the ETS binding sites in the *ins-1* promoter are necessary for the functional activity of *ins-1 in vivo.* Therefore, we used the mutated *ins-1* promoter to drive *ins-1* cDNA (*ins-1pΔΔETS∷ins-1*), and tested whether this construct could rescue the *ins-1(nj32)* exploration and fat phenotypes. We found that this construct failed to rescue the *ins-1(nj32)* exploration defect in three independent transgenic lines (Figure 4F), in contrast to the non-mutated promoter, which significantly rescued in all lines tested (Figure 3B). We also found that mutating the ETS sites within the *ins-1* promoter region abolishes the ability of this construct to rescue *ins-1(nj32)* fat levels (Figure 4G-H). These data show that the ETS sites in the *ins-1* promoter are required for correct spatial regulation of *ins-1* to enable correct *ins-1* gene function.

To examine if ETS-5 can directly bind the *ins-1* promoter *in vivo*, we used CRISPR/Cas9 to generate a strain where ETS-5 is C-terminally GFP-tagged at the endogenous locus (ETS-5-GFP) (Figure 5). ETS-5-GFP is expressed in the BAG neurons and other head neurons, including the ASG and OLQ neurons, confirming previous reporter-gene analysis and recent single-cell RNA-seq analysis (Figure 5C, (Brandt et al., 2012; Cao et al., 2017; Guillermin et al., 2011)). Importantly, the endogenous expression levels of ETS-5 co-localise in the BAG and ASG neurons with activity of the *ets-5* promoter previously shown to rescue the ETS-5 exploration defect (Figure 5D) (Juozaityte et al., 2017). This engineered strain displays wild-type exploration, suggesting that addition of GFP does not disrupt ETS-5 function (Figure 5E). We used this engineered strain to perform Chromatin Immunoprecipitation with quantitative PCR (ChIP-qPCR), using an α-GFP antibody, to determine if ETS-5 directly binds to the *ins-1* promoter. Our data show significantly higher ChIP enrichment of the *ins-1* promoter region that contains the ETS sites, compared to an upstream control region (Figure 5F). These data demonstrate that ETS-5 binds to the *ins-1* promoter to directly regulate *ins-1* expression.

**Figure 5.**
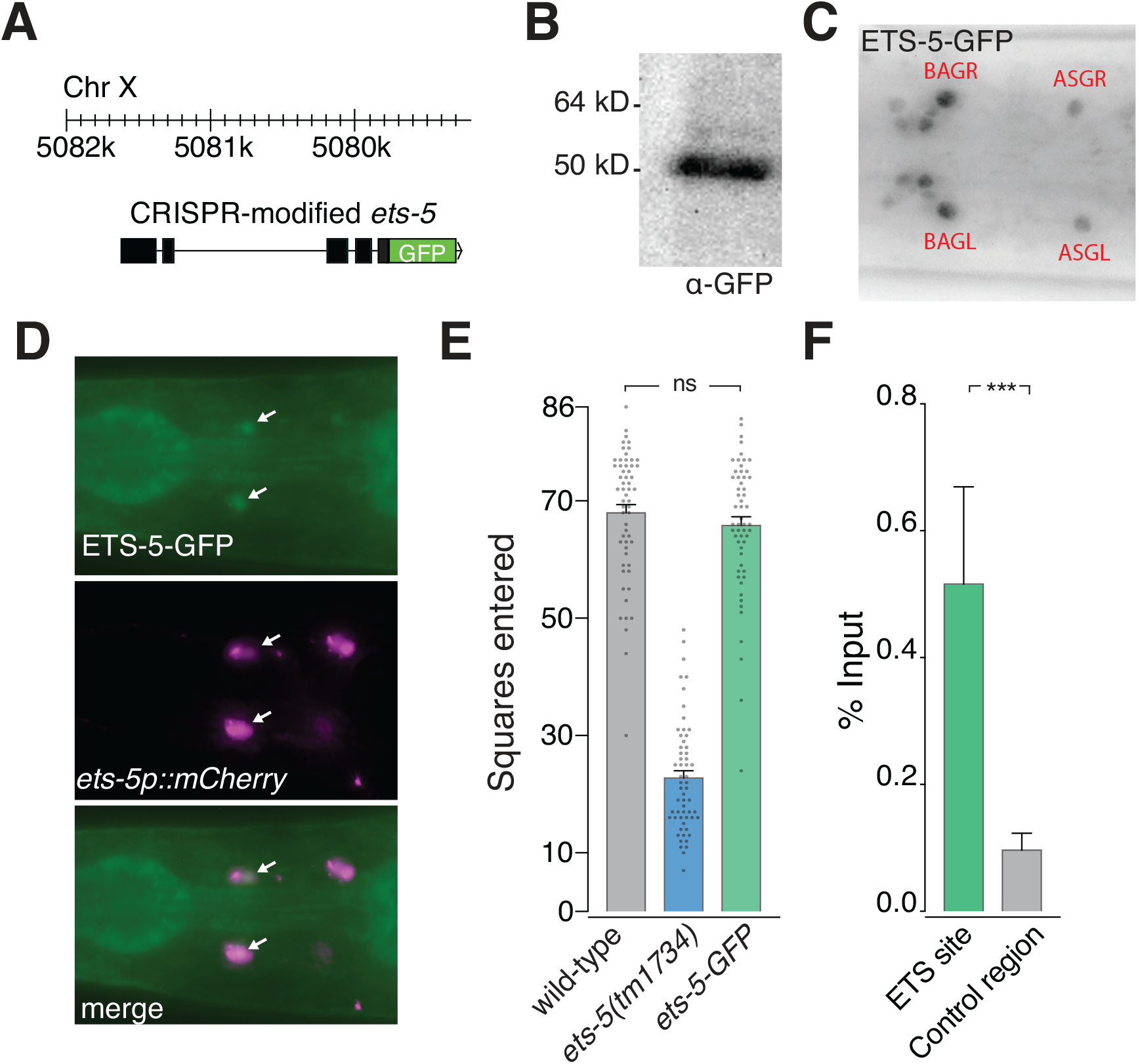
Endogenously tagged ETS-5 binds the *ins-1* promoter. (A) Schematic showing location and construction of C-terminal GFP tagged ETS-5. (B) ETS-5-GFP western blot analysis. ETS-5 (22.6kDA) + GFP (~27kDa) = ~50kDa. (C) Endogenous ETS-5-GFP is expressed in multiple neurons, including BAG. (D) Endogenous ETS-5 expression and functionally sufficient promoter region colocalise in BAG neurons (red arrows) and ASG neurons (green arrow). (E) Exploration assay showing ETS-5-GFP explores to the same extent as wild-type animals. Data presented as individual worm (points) with 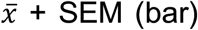, n>50. ns – not significant (One-way ANOVA with Tukey’s correction). (F) ETS-5-GFP ChIP-qPCR. Data are presented as 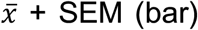, n = 5. **p<0.01 (ratio paired t test.)

### ETS-5 and INS-1 Dynamically Respond to Changes in Diet

We have shown that the ETS-5 transcription factor directly controls BAG-specific expression of *ins-1*. We hypothesized that this neuron-specific gene regulatory mechanism may be important for coordinating responses to ephemeral habitats and nutrient availability. To determine if *ets-5* or *ins-1* expression is dynamic in response to altered nutritional status, we incubated worms on *E. coli* OP50 exposed to a high glucose diet (HGD = 40mM glucose) for 24 hours. Worms fed OP50 grown on glucose have increased triacylglycerides, but carbohydrate levels are unaffected (Brooks et al., 2009). We, and others, have shown that feeding worms on OP50 + glucose drastically increases intestinal fat levels (Figure 3F, (Garcia et al., 2015; Juozaityte et al., 2017)). We postulated that this increase in energy availability may elicit a response of the ETS-5/INS-1 regulatory module in the BAG neurons. Indeed, after 24 hours of feeding on a HGD, the level of endogenous ETS-5-GFP in the BAG neurons significantly decreases (~20% reduction on HGD) compared to OP50-fed control worms (Figure 6A-B). We repeated the same experiment, using the *ins-1p∷NLS-GFP* reporter line and found that *ins-1* promoter activity dramatically decreased on high glucose diet (~70% reduction on HGD, Figure 6C-D). Calculated total fluorescence of the ASH control neuron did not significantly change between the control and glucose conditions (Figure S4). These data suggest a mechanism where, in situations where energy availability is high, nuclear ETS-5 levels are reduced, resulting in a decreased level of binding and activation of the *ins-1* promoter (Figure 6E). We propose that this leads to a decrease in INS-1 expression within the BAG neurons. Reducing INS-1 signalling from the BAG neurons would have the effect of promoting fat storage and enhancing quiescence behaviour.

**Figure 6.**
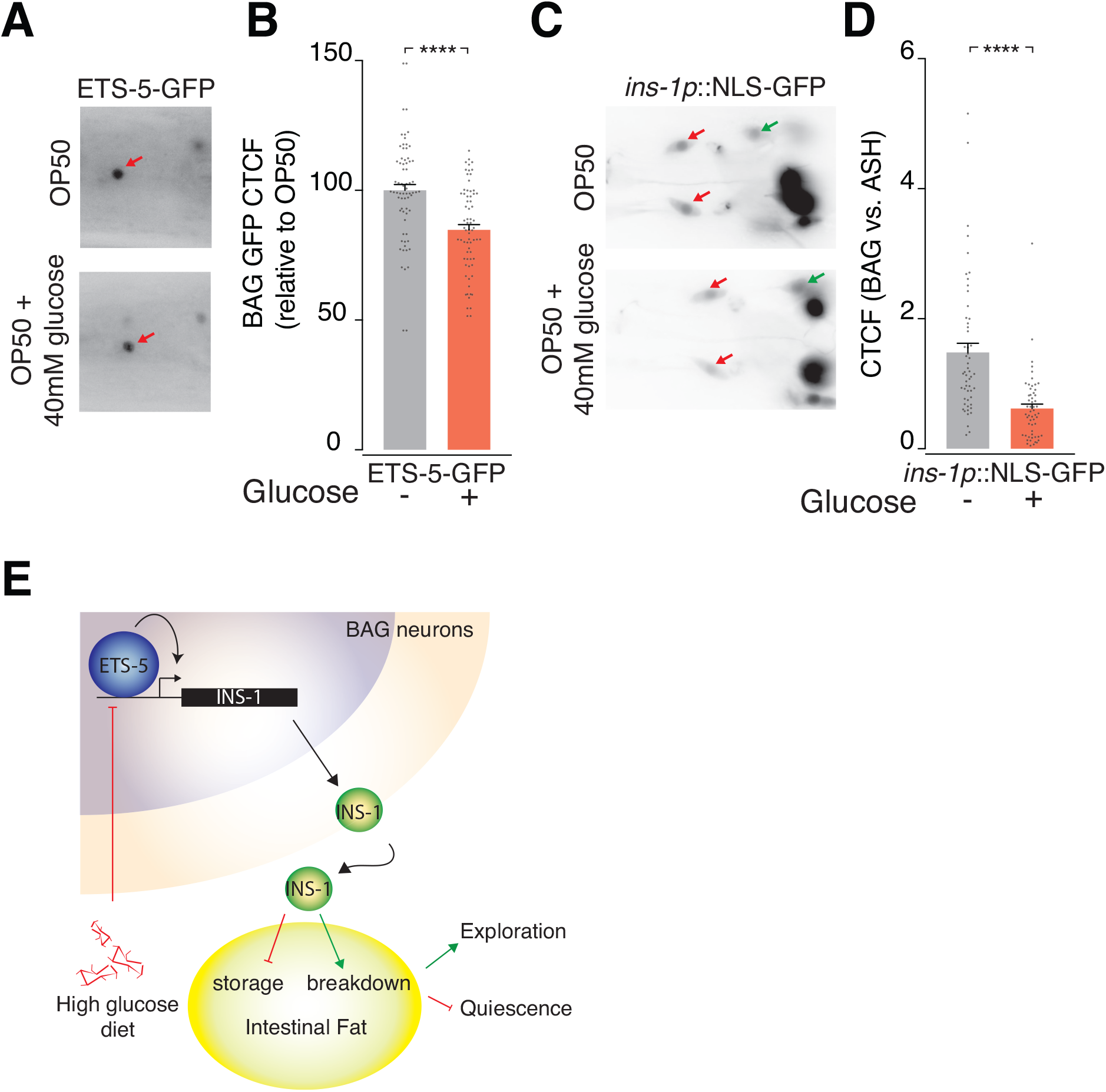
ETS-5 and INS-1 dynamically respond to changes in nutrient availability. (A) Representative images of ETS-5-GFP intensity in the BAG neurons (red arrow) of OP50-treated (upper panel) and OP50 + 40mM glucose-treated (lower panel) animals. (B) Quantification of ETS-5-GFP in the BAG neurons. Treatment: 24 hours. Data presented as individual nuclei measurements (points) with 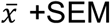, n=70. ****p<0.0001(unpaired t test). (C) Representative images of *ins-1p∷NLS-GFP* expression in OP50-treated (upper panel) and OP50 + 40mM glucose-treated (lower panel) animals (BAG neurons = red arrows, ASH = green arrow). (D) Quantification of GFP intensity in the BAG neurons relative to ASH control neurons in the *ins-1p∷NLS-GFP* reporter grown on OP50 or OP50 + 40mM Glucose. Treatment 24 hours. Data presented as individual nuclei measurements (points) with 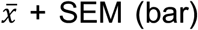, n>50. ****p<0.0001(unpaired t test). (E) Model of the ETS-5/INS-1 regulatory module. Within the BAG neurons, ETS-5 binds to and regulates *ins-1* expression. INS-1 acts from the BAG neurons to inhibit fat storage or promote fat breakdown in the intestine, which induces changes to foraging behaviour. Worms fed a high glucose diet down-regulate ETS-5 and INS-1 levels in the BAG neurons. This suppression of ETS-5/INS-1 action would promote fat storage and quiescence behaviour.

## DISCUSSION

Here we have shown that ETS-5 is not only important for establishing the identity of the BAG neurons, but continues to shape the functional output of BAG neurons after they are formed. ETS-5 facilitates this plasticity by regulating the expression of an insulin-like peptide, INS-1, when animals are grown on a glucose-supplemented food source. We found that INS-1 acting from the BAG neurons, but not from other INS-1-producing neurons, controls intestinal lipid stores, which in turn modulate exploration behaviour.

Although the gas-sensing ability of BAG neurons is not necessary for controlling exploration behaviour, this sensory function of BAGs should not be ignored. Low O_2_ can activate the BAG neurons, which then act to supress the URX neurons through the neuropeptide FLP-17. Under low oxygen conditions, FLP-17 inhibition of URX modulates intestinal lipid breakdown during starvation (Hussey et al., 2018). However, we found that *flp-17* mutant animals have wild-type levels of exploration (Figure 1B). These observations indicate that ETS-5/INS-1 act in a separate regulatory module within the BAG neurons from O_2_/FLP-17. These BAG-specific mechanisms likely act in parallel to ensure appropriate metabolic regulation depending on the environmental context of the animal. Since ETS-5 regulates expression of the FLP-17 and FLP-19 neuropeptides (Brandt et al., 2012; Guillermin et al., 2011), O_2_ regulates FLP-17 levels (Hussey et al., 2018), and CO_2_-sensing is required for FLP-19 expression (Rojo Romanos et al., 2017), these regulatory modules are likely to be linked in a broader physiological sense – suggesting the idea that activity within the BAG neuron can be precisely tailored according to multiple sensory inputs.

INS-1 activity from the BAG neurons itself appears to be of a highly precise nature. We observed that the *ins-1* promoter containing mutated ETS-binding sites was still active in non-BAG *ins-1* expressing neurons, such as the ASH and AIA (Pierce et al., 2001). This means that *ins-1* is still expressed in these other INS-1-positive neurons in the *ins-1pΔΔETS∷ins-1cDNA* transgenic lines, yet fails to rescue the *ins-1(nj32)* fat and exploration phenotypes. Therefore, our data suggest that BAG-produced INS-1 possesses specific properties that differentiate it from non-BAG-derived INS-1 that enable it to perform its function in controlling fat metabolism. Our data clearly show that simply expressing *ins-1* anywhere in the nervous system is not sufficient for INS-1 to control intestinal metabolism. It has been previously shown that INS-1 acting from the AIA interneuron can control odour-evoked responses in AWC neurons (Chalasani et al., 2010), and regulate salt-chemotaxis learning through DAF-2 in the ASE neurons (Tomioka et al., 2006). In these two AIA-specific examples, INS-1 controls sensory neurons that the AIA synapses to, demonstrating that the site of INS-1 action can be highly specific and localised. In our study, INS-1 signals from a sensory neuron, therefore the direction of action is different than the above examples. The BAG neurons chemically synapse onto multiple interneurons: RIA, RIB, AIY, AIB, AVE, RIH, RIG, AIN and AVA (White et al., 1986). Therefore, it could be possible that BAG-specific INS-1 signals to one (or more) of these interneurons, which in turn regulates fat levels and exploration behaviour. BAG-specific INS-1 could also act on other neuron types such as the URX, which is responsive to BAG-expressed FLP-17 activity. Alternatively, as INS-1 is secreted via dense-core vesicles into the pseudocoelomic fluid (Laurent et al., 2018; Tomioka et al., 2006), it is possible that the specificity of BAG-INS-1 lies not in spatial positioning, but instead may be achieved through timing of release, or post-translational modifications.

Active neuropeptides are produced by processing pro-neuropeptide proteins. Recent progress in mass spectrometry analyses is revealing the highly complex nature of pro-neuropeptide modification (reviewed in (DeLaney et al., 2018; Van Bael et al., 2018b)). A single pro-neuropeptide may be processed into multiple neuropeptides with differing activities, thereby extensively expanding *C. elegans* signalling capacity (Bargmann, 2012; Nassel, 2009). Pro-neuropeptide processing includes peptide cleavage, such as removal of the c-peptide in INS-1, and formation of disulphide bonds. In addition to this, neuropeptides may undergo potential modifications such as acetylation, phosphorylation, glycosylation, and amidation (Van Bael et al., 2018a), which may change the bioactivity of the neuropeptide (reviewed in (De Haes et al., 2015)). For example, Leinwald and Chalasani (Leinwand and Chalasani, 2014) suggest that the processing of the INS-6 pro-peptide by different enzymes may lead to INS-6 becoming amidated in the ASI neurons, but not in ASE neurons – which could generate INS-6 neuropeptides with different stabilities and modes of action from the same INS-6 pro-peptide (Hung et al., 2013; Leinwand and Chalasani, 2013). Differential processing of the INS-1 proprotein, in combination with spatial and temporal regulation of INS-1 release, could ensure BAG-INS-1 acts specifically and appropriately to regulate metabolism.

Our analysis of whether *ins-1* acts through the canonical insulin/IGF-like receptor DAF-2 yielded conflicting results that can be most simply attributed to the differing sensitivities of the two assays used – Oil Red O fat staining being more sensitive than the exploration assay. Our fat staining data show the *daf-2; ins-1* double mutant stored significantly more fat than the *daf-2* single mutant. However, the impact that this additional fat storage would have on exploration behaviour can only ever be minimal, as the *daf-2* mutant exploration phenotype is already very near the minimum threshold of our assay 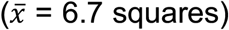. Therefore, we are unlikely to detect if the *daf-2; ins-1* double mutant exhibits an additive effect on exploration. With the Oil Red O analysis, we observed that DAF-2 is likely not required for the role of INS-1 in regulating fat storage. As the *daf-2(e1370)* allele is not a null, but a temperature-sensitive missense mutation, we cannot rule out completely that *ins-1* is not acting through *daf-2*. However, our data showing that *daf-16* has no effect on exploration, and does not alter the exploration phenotype of either *ets-5(tm1734)* or *ins-1(nj32)*, further supports a DAF-2-independent role of BAG-INS-1. That DAF-2 is not required for BAG-specific INS-1 activity supports the presence of alternative receptors for insulin-like molecules in *C. elegans* (Chalasani et al., 2010). Identifying these alternative receptors, where they act, and their downstream targets, are crucial future avenues of investigation for understanding this neuron-intestinal communication paradigm.

The mechanism of ETS-5 regulating *ins-1* expression in *C. elegans* is likely to be conserved in mammals - as Pet1/Fev, the mammalian orthologue of ETS-5, regulates insulin expression in the pancreatic β-cells (Ohta et al., 2011). However, the biological outcomes of *C. elegans* INS-1 and mammalian insulin are opposite: INS-1 promotes fat mobilisation and is downregulated in high glucose conditions, whereas mammalian insulin is secreted in high glucose conditions, and promotes glucose uptake and fat storage (reviewed in (Dimitriadis et al., 2011)). While it has been previously shown that feeding human insulin to *C. elegans* ameliorates the negative effect of a high glucose diet on lifespan (Mendler et al., 2015), and that human insulin can antagonise DAF-2 to promote dauer arrest (Pierce et al., 2001). Here, we found that human insulin cannot functionally replace *ins-1* in BAG neurons (Figure S1C), further supporting the idea that BAG-derived INS-1 acts in a specific, rather than general, context to regulate metabolism. Like ETS-5, Pet1 is also expressed in the nervous system, where it functions to specify serotonergic neuron fate, although there are non-serotonergic Pet1 positive domains in the central nervous system (Hendricks et al., 1999; Hendricks et al., 2003; Pelosi et al., 2014). It is therefore possible that Pet1 regulates an insulin-related peptide, such as a relaxin, in mammalian neurons in a similar way as in *C. elegans*.

We found that endogenous ETS-5 levels are decreased in the BAG nucleus after animals are grown for 24 hours on a fat-inducing diet. Under the same conditions, there is a more substantial decrease in *ins-1* promoter activity observed, which we attribute to decreased ETS-5 binding at the *ins-1* promoter. ETS-5 levels are significantly reduced after 24 hours of feeding on a fat-inducing diet, indicating that the effect of diet on BAG function occurs on a vastly slower timescale than O_2_/CO_2_ responses in these neurons - seconds for responding to changes in O_2_ (Skora and Zimmer, 2013; Zimmer et al., 2009). Determining whether the decrease in ETS-5 levels is a result of shuttling the transcription factor out of the nucleus, increased protein degradation, or decreased translation are important future questions, which will enable us to understand how this mechanism is controlled by dietary changes. One consequence of ETS-5 downregulation under fat-inducing conditions could be that other BAG-specific functions are also altered. The expression of the guanylate cyclases required for O_2_ and CO_2_ sensing, and the neuropeptides that elicit the downstream signals from BAG activation through gas sensing, are all dependent on ETS-5 to some extent (Brandt et al., 2012; Guillermin et al., 2011). Therefore, the decrease in ETS-5 within the BAG neurons under high-nutrient conditions could broadly impact the sensing and signalling functions of these neurons, which would have implications for the physiology of the entire animal – from behaviour, to egg laying, to intestinal metabolism.

## ACKNOWLEDGEMENTS

We thank Mark Febbraio, Corey Laverty and members of the Pocock Laboratory and for advice and comments on the manuscript. Some strains were provided by the *Caenorhabditis* Genetics Center (University of Minnesota), which is funded by NIH Office of Research Infrastructure Programs (P40 OD010440), and by the National BioResource Project of Japan. We extend our thanks to Chris Hopkins and Trisha Brock (Knudra Transgenics) for their expertise in genome-editing to generate CRISPR/Cas9 tagged ETS-5-GFP. This work was supported by the following grants: NHMRC (Projects GNT1105374 and GNT1137645 to R.P.) and veski Innovation Fellowship (VIF23 to R.P.).

## AUTHOR CONTRIBUTIONS

Conceptualization, A.H. and R.P.; Methodology, A.H., Q.W., T.S., and R.P.; Investigation, A.H., Q.W., T.S., and R.P.; Writing – Original Draft, A.H.; Writing – Review & Editing, A.H., Q.W., T.S., and R.P.; Funding Acquisition, R.P.; Resources, R.P; Supervision, A.H. and R.P.

## STAR Methods

### CONTACT FOR REAGENT AND RESOURCE SHARING

Further information and requests for resources and reagents should be directed to and will be fulfilled by the Lead Contact, Roger Pocock.

### EXPERIMENTAL MODEL AND SUBJECT DETAILS

#### Mutant and transgenic reporter strain

Strains were grown using standard growth conditions on NGM agar at 20°C on *Escherichia coli* OP50, unless otherwise stated (Brenner, 1974). All mutant strains were backcrossed to N2 a minimum of 3 times prior to experiements.

#### Transgenic line

All constructs were injected into young adult hermaphrodites as complex arrays. *ins-1* rescue constructs were injected into *ins-1(nj32)* background: 2 ng/μl of construct (fosmid *WRM0618bG05*, *ins-1p∷ins-1∷SL2∷GFP*, *ins-1pΔETS1ΔETS2∷ins-1∷SL2∷GFP, flp-17p∷ins-1∷SL2∷GFP*) were injected with 5 ng/μl of *myo-2p∷mCherry* and 180 ng/μl bacterial DNA. While the GFP of the rescue construct was visible in some cases, the rescuing concentrations used were too low to reliably see GFP expression. *ins-1* promoter reporter assays: 50 ng/μl of construct (*ins-1p∷NLS∷GFP*, *ins-1pΔETS1∷NLS∷GFP, ins-1pΔETS2∷NLS∷GFP, ins-1pΔΔETS∷NLS∷GFP)* injected with 50 ng/μl *rol-6* co-injection marker and 180 ng/μl bacterial DNA. Microinjections were performed using standard methods (Mello et al., 1991).

#### CRISPR/Cas9

The endogenously tagged ETS-5∷GFP strain was generated by Knudra Transgenics (now NemaMetrix). Genomic modifications include: Last 34 amno acids of ETS-5 are recoded with silent mutations to prevent recutting. GGGGSYG linker with intron inserted, and codon-optimised eGFP inserted C-terminally. TGTGSGSSTS linker with intron containing the loxp flanked unc-119 rescue cassette and stop codon.

### METHOD DETAILS

#### Molecular cloning

##### ins-1p reporter construct

A genomic fragment was PCR-amplified from the putative TSS of the *ins-1* gene to 2.574 kb upstream. The forward primer incorporated a HindIII site, and the reverse primer incorporated a BamHI site. The HindIII and BamHI digested promoter fragment ligated into HindIII and BamHI digested pD95.67 vector, resulting in *ins-1p∷NLS∷GFP*

##### Promoterless *ins-1cDNA* construct

To generate a construct containing *ins-1* cDNA that would enable insertion of multiple promoter fragments: *ins-1 cDNA* (330 bp) was amplified from N2-derived cDNA incorporating NheI and KpnI sites. The *ins-1 cDNA* insert was cloned using NheI-KpnI into MCS2 of the pSM_SL2_GFP vector.

##### *ins-1p∷NLS-GFP ETS-site* mutagenesis

The ETS sites within the *ins-1p∷NLS-GFP* reporter construct were mutated using site-directed mutagenesis. First, individual sites were modified independently and sequenced, then the *ins-1pΔETS2∷NLS-GFP* construct was used for site-directed mutagenesis of ETS1 to generate *ins-1pΔΔETS∷ins-1∷SL2∷GFP*.

##### *ins-1p∷ins-1cDNA* rescue construct

The *ins-1* promoter was PCR amplified from the *ins-1p∷NLS-GFP* construct, incorporating SmaI sites to non-directionally clone into MCS1 of the pSM_ins-1cDNA_SL2_GFP vector. Directionality was confirmed by sequencing.

##### *flp-17p∷ins-1cDNA* rescue construct

A 200 bp fragment of the *flp-17* promoter was cloned into the pSM_ins-1cDNA_SL2_GFP by restriction-free cloning (Bond and Naus, 2012). The flp-17 promoter was amplified by PCR from *Prom9flp-17∷mCherry* (Brandt et al., 2012).

##### *flp-17p∷ins-1ΔΔCS* rescue construct

The cleavage sites of *ins-1* within the *flp-17p∷ins-1cDNA* construct were mutated by site-directed mutagenesis.

##### *flp-17p∷human insulin* rescue construct

The Human insulin coding sequence was synthesized by BIONEER PACIFIC, Engineered with NheI and KpnI sites to clone by NheI-KpnI digest into the *flp-17p∷ins-1cDNA_SL2_GFP* vector.

##### *ins-1p∷ins-1cDNA ETS-site* mutagenesis

The *ins-1pΔΔETS∷ins-1cDNA* construct was generated as described for *ins-1pΔΔETS∷ins-1∷SL2∷GFP* above, sequentially mutating ETS2 then ETS1 using *ins-1p∷ins-1cDNA* as a mutagenesis template.

#### Exploration assay

Cultures for exploration assays were maintained in low-population density, non-starved condition for at least two generations prior to commencing experiments. On day one of the assay, five L4 animals were placed on NGM plates freshly seeded with OP50 and cultured at 20°C for four days. In parallel on day one, unseeded NGM plates were taken from 4°C storage and placed at room temperature in stacks of two, to ensure even drying. On day three, a single colony of OP50 from a freshly streaked (within two weeks) plate was used to inoculate 400 mL of LB, which was incubated for 16 hours at 37°C without shaking, then placed at 4°C until required. In the afternoon of day four, each NGM plate was uniformly seeded with 400 μL of the OP50 culture, and left at room temperature until required for the assay. On day five at 9am 25 Christmas-tree L4 larvae from each genotype, were picked to fresh NGM+OP50 plates. At 5pm on day five, single animals were transferred to the centre of a uniformly-seeded plate, placed on the lab bench, and left undisturbed for 16 hours. Twenty worms were tested for each genotype, for each assay, and stacked 2 plates high during the assay. After 16 hours (day six of the experiment) worms were removed from the assay plates and the number of squares the tracks in the OP50 enter was counted (4.05 inches/line, 86 squares maximum). Exploration assays were repeated in biological triplicate on different days, always in parallel with a wild-type control, and blinded prior to counting squares entered.

#### RNAi experiment

RNAi of *pod-2* exploration assay was performed as **exploration assay** above with the following differences: On day dne L4s animals were placed onto RNAi plates uniformly seeded with L4440 (empty-vector) bacteria. RNAi exploration plates were incubated at 37°C for 16 hours, 24 hours before the exploration assay (day four). In parallel, 5 mL bacterial cultures of L4440 (empty-vector) or L4440_*pod-2* RNAi-containing HT115 *E. coli* were grown in LB + Ampicillin. After the 16 hours drying (day 5), the RNAi plates were uniformly seeded with L4440 or *pod-2* RNAi bacteria by coating the plates 2-3 mL of bacterial culture, tipping the culture off, then removing excess culture by flicking the plate sharply 3 times onto paper towel. The plates were then incubated at 37°C for 7 hours, then moved to room temperature for one hour before beginning the exploration assay. Christmas-tree L4s were picked to the required RNAi plates (made previously) 8 hours before beginning the exploration assay, resulting in animals being exposed to the RNAi for a total of 24 hours for the assay.

#### Oil Red O staining

Oil Red O staining was performed essentially as described previously (Noble et al., 2013):

##### Worm synchronisation

Five L4s were picked to four fresh NGM+OP50 plates per genotype, and allowed to grow for 5 days. Worms were washed from the plates in 4 mL of M9 buffer into a 15 mL conical tube (Thermo Scientific) and eggs were isolated by adding 0.5 mL bleach (White King Premium)+ 0.5 mL 5 M NaOH, mixing and incubating for 4 minutes. M9 was added to fill the tube, then samples were centrifuged for 1 minute at 1000 RCF, and the supernatant removed. Egg pellets were washed a further 3 times in M9 buffer, centrifuging for 1 minute at 1000 RCF. The egg pellet was resuspended in ~1 mL M9 and passed through a 40 μm filter to a fresh 15 mL tube. Samples were kept at room temperature, with gentle rocking for 24 hours to allow all worms to hatch and reach the L1 stage. Approximately 600 L1s were plated onto fresh NGM+OP50 plates (~8 plates per genotype), and incubated at 20°C for 64 hours to reach young adulthood.

##### Staining

Oil Red O stock solution was prepared by adding 0.5 g Oil Red O with 100 mL isopropanol. Solution covered with foil and mixed at room temperature for 2 days. Immediately prior to commencing fat staining, Oil Red O stock solution was diluted to 60% with sterile milli-Q filtered water, covered with foil and rotated at room temperature until required. Worms were washed from plates with 10 mL PBS and placed into a 15 mL conical tube using a glass Pasteur pipette, and centrifuged at 3300 RCF for 30 seconds. Supernatant was aspirated to 1 mL volume, worms were resuspended and transferred, with Pasteur pipette, to a 1.5 mL microcentrifuge tube. Worms were centrifuged at 3300 RCF for 30 seconds, supernatant removed to 100 μl volume, and pellet resuspended in 1mL PBS with a brief, gentle vortex to mix. Centrifugation and PBS wash steps were repeated a total of 2 times, finishing with an additional of 1 mL PBS and centrifugation. The samples were incubated on ice for 10 minutes, then supernatant was aspirated to 0.1 mL, 50 μL PBS, and 150 μL fixation buffer (30 mM PIPES, 14 mM Na_4_EGTA, 40 mM NaCl, 160 mM KCl, 1 mM Spermidine, 0.4 mM Spermine, 4 % Formaldehyde, 0.5 % β-mercaptoethanol). Samples were fixed for 5 minutes at room temperature, mixing gently during fixation. Placing tubes in a floating rack, the tubes were placed in a dry ice + ethanol bath for 3 minutes, ensuring contents of tubes were completely frozen. Samples were then partially thawed by placing at room temperature water for 1 minute. Samples were immersed in the dry ice + ethanol bath for 2 minutes, thawed in room temperature water for 1 minute, and again immersed in the dry ice + ethanol bath for 2 minutes. Samples were thawed at room temperature for 45 seconds, then tubes were quickly, and with moderate pressure, swiped across a tube rack to obtain an ice slurry. Tubes were laid down at room temperature until the ice slurry completely melted (5-10 minutes). Samples were centrifuged 3300 RCF for 30 seconds, supernatant aspirated to 100 μl, 1 mL PBS added then briefly vortexed. Wash steps were repeated for a total of three washes. After removing the final supernatant to 100 μl volume, 1mL of 60 % isopropanol was added to the worm pellet, and samples were incubated for 10 minutes with rotation at room temperature. Samples were centrifuged 3300 RCF for 30 seconds and maximum supernatant was removed without disturbing the worm pellet. Oil Red O working solution was filtered through a 0.22 μm filter, then 400 μl was added to each worm pellet. Samples were covered in foil, then incubated with rotation, overnight at room temperature. The following day, the stained worms were washed twice in PBS + 0.01% Triton-X100, then washed once in PBS, centrifuging 3300 RCF 30 seconds. Supernatant was removed, then worms were resuspended and mounted onto agarose (as below without NaN_3_) pads for imaging.

#### Fluorescence microscopy

Animals were anesthetized with 20 mM NaN_3_ on 5% agarose pads, and images were obtained with an Axio Imager M2 fluorescence microscope, Axiocam 506 mono camera and Zen software (Zeiss). Fluorescence images were obtained using a 100 X oil objective, and collected as Z-stacks through the worm head with 1 μm step size. Oil Red O images were obtained with a 40 X objective, using mCherry, GFP, and DAPI filters with transmitted light to obtain RGB images.

#### Chromatin Immunoprecipitation – qPCR

##### Sample preparation

ChIP-qPCR of ETS-5∷GFP was performed on non-starved synchronised L4 animals collected as follows. On day one, approximately 100 L4 hermaphrodites were picked to fresh NGM+OP50 plates. Day two, 15 animals (now young adult) were transferred to fresh NGM+OP50 plates and allowed to lay eggs for four hours, then removed. Eggs were incubated at 20°C for 67 hours (~10 eggs/adult stage). Animals were washed from NGM plates with M9 buffer, and incubated with 2.5 mL bleach and 2.5 mL 5 M NaOH for four minutes to release eggs. Eggs were then washed 3-4 times in M9 buffer, centrifuging at 1000 RCF, 1 minute between wash steps. Eggs were plated at a density of ~600/plate to fresh NGM+OP50 plates. 62-63 hours after egg plating, late L4s were washed off plates with M9, centrifuged at 1000 RCF for 1 minute, and washed two times in M9. L4s were then washed once in PBS + protease-inhibitor complex (PIC). L4s were then resuspended in 1mL PBS + PIC and snap frozen as droplets in liquid nitrogen (“popcorn”) that were stored at−80 °C until required.

##### Chromatin preparation

To prepare chromatin: frozen popcorn was ground with a mortar and pestle on dry ice to a fine powder. The frozen powder was transferred to a glass Dounce homogenizer containing 20 mL PBS + PIC + 1.1% formaldehyde. Samples were cross-linked at room temperature for 10 minutes - during this time the samples were ground using the tight “B” pestle. 1 mL of 2.5 M glycine was added to the sample to quench the formaldehyde for 5 minutes at room temperature, while still homogenizing the sample. Samples were then centrifuged for 5 minutes at 6000 RCF, the supernatant removed and pellet washed once with PBS+PIC. The centrifugation step was repeated, supernatant removed and the pellet resuspended in 500 μl FA buffer (50 mM HEPES, pH 7.5, 1 mM EDTA, 1% Triton-X100, 0.1% Sodium deoxycholate, 150 mM NaCl, PIC). Samples were sonicated using a Covaris S220 sonicator in 150 μl tubes using the following settings: Duty cycle 2 %, Intensity 6, cycles/burst 200, for 12 minutes. After sonication of all samples was complete, they were centrifuged at full-speed for one minute (room temperature), the supernatant was kept on ice and the pellet resuspended in 150 μl FA buffer and sonicated an additional time under the same conditions. All sonicated samples were pooled, centrifuged at maximum speed at 4°C for 10 minutes. The supernatant was collected, DNA concentration measured using a nanodrop, and frozen at−80°C until required for Chromatin immunoprecipitation (ChIP).

#### Chromatin Immunoprecipitation

20 μg of chromatin was made to 650 μl volume with FA buffer in a low bind microcentrifuge tube (Eppendorf). Samples were incubated with 2 μl of α-GFP antibody (ab290 abcam) with overnight rotation at 4°C. For each sample, 10 μl Protein-G dynabeads (Invitrogen) was added to a fresh low bind tube, and washed once in 1 mL FA buffer. 500 μl chromatin/antibody sample was added to the beads, and 50 μl chromatin/antibody sample was kept in a fresh low bind tube as input (10%). Samples were incubated with beads for 3 hours, with rotation at 4 °C. Beads were then washed 2 times with 1 mL FA buffer, with 5-minute rotation at 4 °C. Then washed first with 1 mL FA-1M buffer (FA buffer with 1 M NaCl), then LiCl buffer (250 mM LiCl, 1% NP-40, 1% sodium deoxycholate, 1 mM EDTA, 10 mM Tris-HCl, pH 8) for 10 minutes with rotation at 4°C. Finally, beads were washed twice, quickly with TE Buffer then resuspended in 100 μl TE. 50 μl TE was added to input samples. Tubes were then wrapped in parafilm, and incubated overnight at 65°C with 1000 rpm shaking.

##### DNA purification

1 μl RNAseA (Qiagen) was added to each sample, then incubated at 37°C for 30 minutes with shaking. Then, 5 μl 10% SDS and 1 μl 10 mg/mL Proteinase-K solution was added to each sample, and incubated 55°C for 1.5 hours with 1000 rpm shaking. DNA was then purified using the Qiagen min-elute kit as follows: 600 μl PB was added to each sample, mixed thoroughly by pipetting. Tubes were then placed on a magnetic rack to clear beads from solution. All sample was then added to mini-elute columns, centrifuged 1300 rpm 1 min. Columns were washed 2x with 600 μl PE buffer, then columns dried by a further 2-minute centrifugation. 20 μl EB buffer was added to the column, left to stand for 2 minutes, then centrifuged 1 minute to elute the DNA.

##### qPCR analysis

ChIP and input samples were diluted 1:10 in PCR-grade water (Roche), and 4 μl of each diluted sample was pipetted in triplicate and mixed with 1 μl 10 μM primer mixture for control region or *ins-1* promoter ETS region, and 5 μl SYBR green (Roche) and analysed using the Light Cycler 480 (Roche).

#### Western Blot

Protein gels were made using the mini-PROTEAN system (BioRad). Eight non-synchronised, non-starved plates of worms were washed off in M9. Centrifuged at 1700 rpm, washed 3 times in M9, and the supernatant removed. Samples were transferred to 1.5 mL micro-centrifuge tubes, then 50μl loading buffer was added and samples boiled for 5 minutes. Samples were centrifuged and full amount was loaded into the wells of a 10 % Tris-Cl acrylamide gel, including 10 μl protein standard ladder (Novex). Gels were run for 45 minutes at constant mAmps of 0.04. Proteins were transferred to PVDF membranes using iBlot ministacks (Invitrogen) and iBlot device (Life technologies), 20 V 7 minutes. Membranes were blocked in TBST (20 mM Tris,120 mM NaCl, 0.05% Tween 20) + 5% skim milk powder, for 1 hour at room temperature. Membrane was then incubated with ab290 1:1000 diluted in TBST+5% skim milk powder for 1 hour. Membrane was then washed 3 x 10 minutes in TBST. The membrane was then incubated at room temperature with HRP-conjugated α-Rabbit IGG 1:5000 in TBST + 5% skim milk powder. Membrane wash steps were repeated at previously, then the membrane was incubated with ECL western blotting substrate (Pierce/Thermo Scientific) solution for 2 minutes and imaged using ChemiDoc MP imaging system (Biorad).

#### High glucose feeding

To prepare glucose-enriched OP50 plates: NGM plates (6 cm/10 mL) were fully covered with 400 μL of glucose (D-(+)-glucose; Sigma-Aldrich) of a 1 M stock solution prepared in diH2O, to reach the desired concentration of 40 mM, and were allowed to dry for 24 h. The next day, 400 μL of *E. coli* OP50 was added on the top of the glucose to cover the entire plate and left for two nights at room temperature. BAG expression level analysis: Worm cultures were grown as for exploration assay, on day five, 25 Christmas-tree L4s for each condition assayed were first transferred to an unseeded NGM plate, then moved to either OP50 control plates or the glucose enriched plates. Worms were incubated at 20 °C for 24 hours before imaging.

### QUANTIFICATION AND STATISTICAL ANALYSIS

All experiments were performed in three independent replicates, n values are indicated in the figure legends of corresponding experiments. BAG fluorescence intensity was quantified in FIJI (ImageJ) by tracing the BAG nuclei from DIC images, then changing to the fluorescence (GFP) channel and the integrated density, mean grey, and area were measured. ASH nuclei were traced through the fluorescence channel. For each measurement a background grey value measurement was made outside of the worm area in order to calculate total cell fluorescence (CTCF). Oil red O staining was quantified in FIJI (ImageJ) by tracing the first 4 intestinal cells proximal to the pharynx. The intensity in these cells was measured in the inverted green channel (where Oil Red O absorbs the light), collecting area, mean grey value and integrated density data. The pharynx region was used as background measurement for Oil Red O staining, and used to calculate the CTCF. CTCF is calculated as: *Integrated Density – (area * mean gray of background)*. ChIP-qPCR Ct values were converted into % input with the following calculation: Calculated 100 % input = Average Ct value (input) – 3.3. Calculated % input = 100*2^(100% input – Ct (ChIP). Statistical analysis was performed in GraphPad Prism 8 using one-way analysis of variance (ANOVA) for comparison followed by Dunnett’s Multiple Comparison Test or Tukey’s Multiple Comparison Test, or unpaired t test, or ratio-paired t test (ChIP-qPCR), indicated in figure legends. Values are expressed as mean ± s.e.m, where possible individual data points have been plotted. Differences with a *P* value <0.05 were considered significant.

