## Supplementary information for "Diet-responsive Transcriptional Regulation of Insulin in a Single Neuron Controls Systemic Metabolism"

**This PDF file includes:**

Figs. S1 to S4

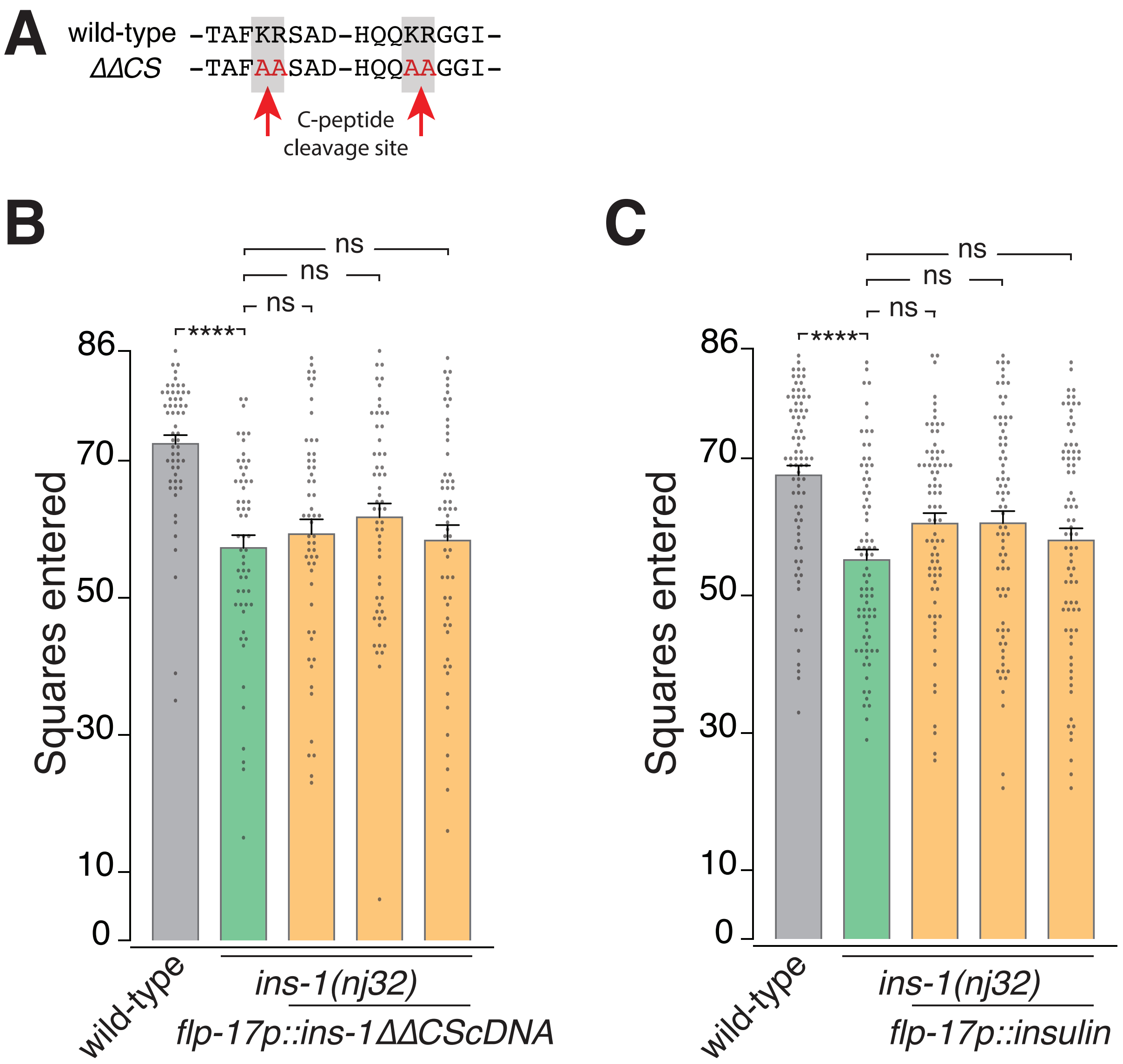


**Figure S1. Non-cleavable INS-1 and human insulin do not rescue *ins-1(nj32)* phenotype from BAG neurons.**

(A) Sequence of mutated ins-1 pro-peptide cleavage sites.

(B) Exploration assay, *ins-1(nj32)* expressing *flp-17p::ins-1cDNA* with mutated cleavage sites (ΔΔCS). Data presented as individual worm (points) with $\bar{x}$ + SEM (bar), n>50. ****p<0.0001, ns – not significant (One-way ANOVA with Tukey's correction).

(C) Exploration assay, *ins-1(nj32)* expressing *flp-17p::insulin* (human insulin). Data presented as individual worm (points) with $\bar{x}$ + SEM (bar), n>50. ****p<0.0001, ns – not significant (One-way ANOVA with Tukey's correction).

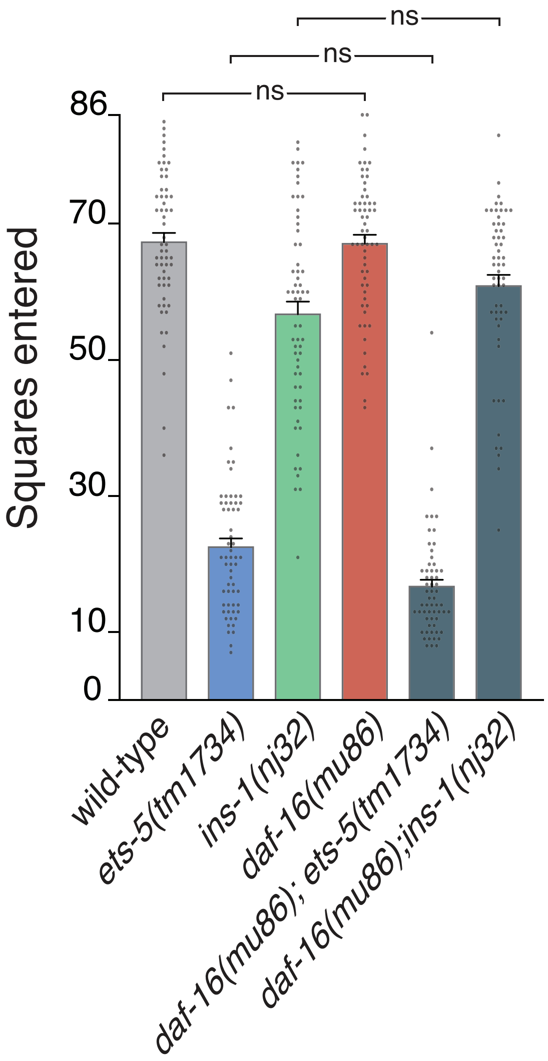


**Figure S2. DAF-16 is not required for ETS-5 or INS-1 control of exploration**

Exploration assay of *daf-16(mu86); ets-5(tm1734)* and *daf-16(mu86); ins-1(nj32)* double mutants. Data presented as individual worm (points) with $\bar{x}$ + SEM (bar), n>50. ns – not significant (One-way ANOVA with Tukey's correction).

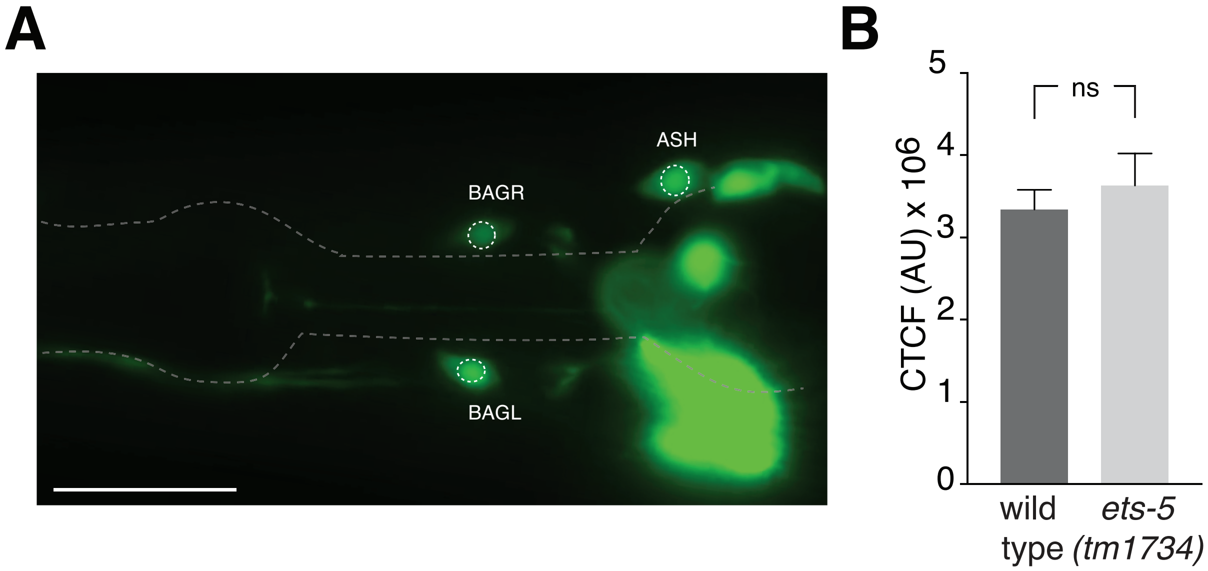


**Figure S3. *ins-1* promoter expression does not change in ASH neurons in *ets-5(tm1734)***

(A) Representative image of *ins-1p::NLS-GFP*, showing measurement regions of both BAG nuclei and a single ASH nucleus. Scale bar = 50µm.

(B) ASH nuclear expression levels of *ins-1p::NLS-GFP* in wild-type and *ets-5(tm1734)* genetic backgrounds. Data are presented as $\bar{x}$ + SEM, n=40 (wild-type), n=39 (*ets-5(tm1734)*). ns – not significant (unpaired t test).

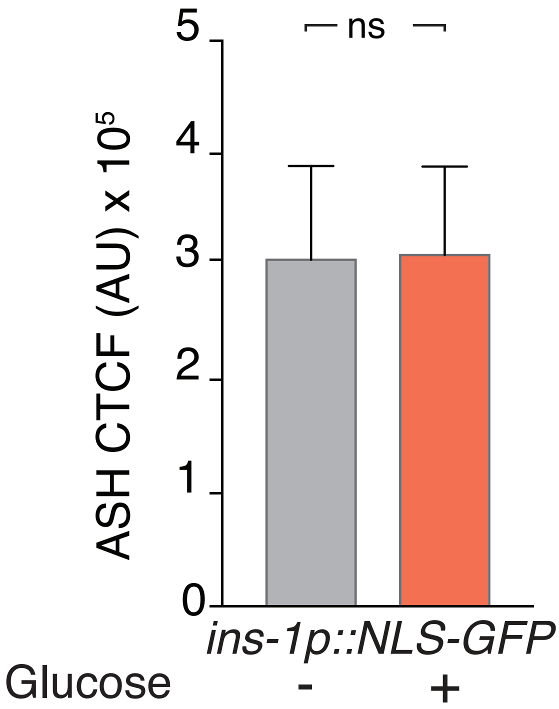


**Figure S4. High glucose diet does not change ASH *ins-1* promoter activity**

ASH nuclear expression levels of *ins-1p::NLS-GFP* do not change under OP50 + 40mM glucose conditions. Data are presented as $\bar{x}$ + SEM, n=27 (OP50), n=28 (Glucose). ns – not significant (unpaired t test).
